# Discrete or indiscrete? Redefining the colour polymorphism of the land snail *Cepaea nemoralis*

**DOI:** 10.1101/383042

**Authors:** Angus Davison, Hannah J. Jackson, Ellis W. Murphy, Tom Reader

## Abstract

Biologists have long tried to describe and name the different phenotypes that make up the exuberant colour polymorphism of the land snail *Cepaea nemoralis*. Traditionally, the view is that the ground colour is one of a few major colour classes, either yellow, pink or brown, but in practise it is frequently difficult to distinguish the colours, and consistently define different shades of the same colour. To understand whether colour variation is continuous, and to investigate how the variation may be perceived by an avian predator, we applied psychophysical models of colour vision to shell reflectance measures. The main finding is that both achromatic and chromatic variation are indiscrete, being continuously distributed over many perceptual units, with the major axis of chromatic variation representing differences in saturation, or purity of colour. Nonetheless, clustering analysis based on the density of the distribution revealed three groups, roughly corresponding to human-perceived yellow, pink and brown shells. There is also large-scale geographic variation between these morphs across Europe, and some covariance between shell colour and banding patterns. Although further studies are necessary to understand the evolutionary origins and impact of natural selective upon this variation, the observation of continuous variation in colour is intriguing, given that the underlying supergene that determines colour should prevent phenotypes from “dissolving” into continuous trait distributions.

---

Throughout the past century, the study of animal colour has been critical in making progress in understanding the principles of biology, especially with respect to genetics and evolution (McKinnon and Pierotti 2010; McLean and Stuart-Fox 2014; Cuthill et al. 2017; San-Jose and Roulin 2017). For instance, early studies on the inheritance of colour traits were important in establishing an understanding of basic Mendelian genetics (Wheldale 1907; Staples-Browne 1908). Subsequently, studies of the distribution and predation of colour morphs have and continue to shape our understanding of how natural and sexual selection operate in wild populations (Hugall and Stuart-Fox 2012; Dale et al. 2015; Delhey et al. 2017). Most recently, candidate gene and latterly genomic approaches have been used to identify the underlying genes that determine the colour differences (references in Hoekstra 2006; McLean and Stuart-Fox 2014; San-Jose and Roulin 2017).

For practical reasons, many of these prior studies have taken advantage of traits that exhibit relatively simple, discrete variation and straightforward inheritance patterns, but this risks missing the extraordinary variation of life forms and colour traits. It is also likely that in nature discrete variation is the exception rather than the rule – and this is becoming more evident as researchers increasingly use instrumentation to measure colour (Montgomerie 2006), rather than being obliged to bin types into human-defined categories.

Historically, two of the most important animals in studying colour polymorphism have been the peppered moth *Biston betularia* and the grove snail *Cepaea nemoralis* and its sister taxon, *C. hortensis* (collectively “*Cepaea*”, the preferred common name), because individuals are relatively easy to collect and study, and the colour morphs show straightforward inheritance. However, while ongoing and long-term studies on these animals continue to provide compelling evidence for the fundamental role of natural selection in promoting and maintaining variation in natural populations, as well as the impact of modern-day habitat change (Silvertown et al. 2011; Cook et al. 2012), progress in understanding the precise mechanism of the polymorphisms has diverged in the two systems.

In the peppered moth, the precise mutation that determines the colour differences was reported to be in a known patterning gene (van’t Hof et al. 2016). In contrast, an understanding of the ‘exuberant’ (Franks and Oxford 2009) polymorphism of *Cepaea* – whether yellow, pink or brown shells and zero to five bands – has stalled since the 1970s. In part, this may be a reaction to the question of Jones *et al.* (1977) on whether the *Cepaea* polymorphism is “a problem with too many solutions?” Actually, the intention of that work was to emphasise the perfect case study provided by *Cepaea*; as simple explanations for phenotypic variation are the exception, Jones *et al.* were making the point that it is important to study organisms for which polymorphism may be explained by a variety of processes, precisely because they are more realistic. Some 40 years later, this comment is still relevant, given that genomic technologies and DNA sequence analyses should allow us to uncover the relative contributions of each of these processes to contemporary diversity – but it is nonetheless understandable that most study systems are still selected because of their relative simplicity.

Despite a general lack of progress, the *Cepaea* polymorphism retains excellent potential as a model system in evolutionary biology. Previous studies have laid the foundations for future progress, including exploiting many long term studies (Cook et al. 1999; Davison and Clarke 2000; Silvertown et al. 2011; Cameron and Cook 2012; Ożgo and Schilthuizen 2012; Cameron et al. 2013; Schilthuizen 2013; Cook 2014; Ozgo et al. 2017), increasing understanding of the pigments and shell proteome (Mann and Jackson 2014; Williams 2017) and especially, using new genomic methods to identify the genes involved (Richards et al. 2013; Kerkvliet et al. 2017). Of particular interest here, we note (as do others, Surmacki et al. 2013) that there is now a pressing need to quantify objectively the polymorphism of *Cepaea* shells, and to understand how this is perceived by an avian predator, because only then can we properly understand how the polymorphism is maintained. The specific problem is that in the past *Cepaea* shell colours have usually been treated as one of three or more distinct classes (e.g. Cain and Sheppard 1954) – yellow, pink or brown – partly due to a lack of objective measures of colour, especially those that can be used in the field and between different observers and contexts (Cain et al. 1960; Cain et al. 1968; Jones et al. 1977). There has been also the significant issue that human perception of colour is not necessarily objective or the same as that of an avian predator (Surmacki et al. 2013).

Now that objective methods of measuring and analysing colour are widely available (Endler 1990; Montgomerie 2006; Maia et al. 2013; Delhey et al. 2015) and easy to use, at least in a controlled laboratory setting, we set out to measure quantitatively the ground colour of *Cepaea* shells, and so to define the nature of the polymorphism. Specifically, by measuring the shell colour of snails collected across the breadth of the European distribution, we used psychophysical models of colour vision to assess how chromatic variation is perceived by birds (but not categorised; Caves et al. 2018).

Previously, Surmacki *et al.* (2013) used quantitative measures of colour on relatively few individuals to assess how shells match to various backgrounds. Here, we investigate the extent to which the distribution of snail shell colour is continuous along the main axes of chromatic variation, using more than a 1000 individuals and Gaussian finite mixture modelling (Scrucca et al. 2016) to test whether colours fall into clusters in multivariate space. We also aimed to understand if quantitative measures on a relatively small sample of shells can describe – rather than explain – geographic patterns in colour morph frequency across Europe, as others have done in much larger qualitative surveys.

The findings have significance for understanding the *Cepaea* polymorphism, and the nature of the selection that acts upon it, as well as more generally highlighting the need to measure colour objectively in other systems, before being able to test for possible explanations.

## Methods

### Data collection

Individual *C. nemoralis* snails were mainly gathered opportunistically by volunteer-led collection and field trips across Europe (Grindon and Davison 2013). Snails were frozen upon arrival at the University of Nottingham, subsequently thawed and the body extracted from the shell. The ground colour and banding of the shell was then scored qualitatively by an experienced person (A.D.) and a student, as either yellow (Y), pink (P) or brown (B), and unbanded (O), mid-banded (M), or all other banding patterns (B, usually five-banded). Subsequent statistical analyses were carried out at the level of the individual and the level of the population (sample site). So that we could compare broad-scale patterns across Europe, larger groups were also used – individual sample sites were therefore grouped into one of six larger groups (Table S1; Figure 1).

**Figure 1.**
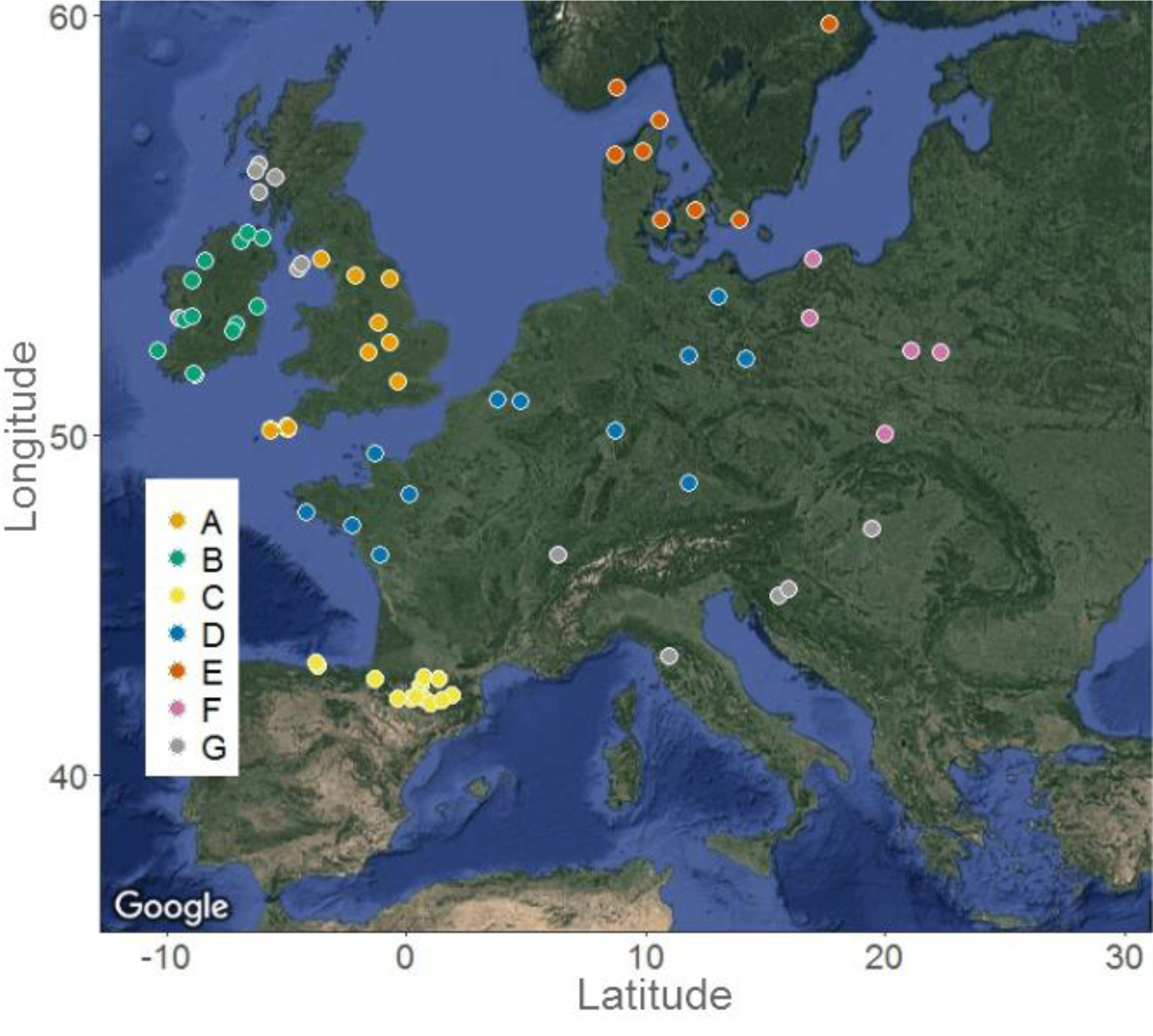
Sample sites across Europe, grouped by geographically contiguous regions. A (England, n=397), B (Ireland, n=144), C (North Spain and Pyrenees, n=112), D (North France, Belgium, Germany, n= 178), E (Scandinavia, n=77), F (Poland, n=126) and all others (n=138).

An Ocean Optics spectrometer (model USB2000+UV-VIS-ES) and a light source (DT-MINI-2-GS UV-VIS-NIR) were used to measure individual reflectance spectra of shells, using a WS-1 diffuse white reflectance standard to set the baseline light spectrum (Teasdale et al. 2013; Taylor et al. 2016), and complete darkness to set the dark spectrum standard. Reflectance measurements were taken on the underside of the dried shell, because it was usually the least damaged region, least exposed to sunlight, and well away from any bands. Point samples were taken for each shell at a 45° incident angle, 2 mm from the shell. Individual shells were measured three times, non-consecutively, with the software recalibrated against light standards every 2-5 measurements. Readings were collected using Ocean Optics SpectraSuite v. 2.0.162 (software settings: integration time 750msec, boscar width 5, scans to average 10); then the raw data smoothed and binned into 5 nm categories using Pavo version 0.5-6 (Maia et al. 2013).

### Analysing chromatic and achromatic variation

We used the framework provided by Delhey et al. (2015) to analyse the reflectance spectra. In this framework, a psychophysical model of colour vision (Vorobyev and Osorio 1998; Vorobyev et al. 1998) is used to assess whether chromatic differences between reflectance spectra exceed a discrimination threshold, or ‘just noticeable difference’ (JND), which can be perceived by a receiver, such as an avian predator. The key to these models lies with the degree to which a particular combination of reflectance and illuminant spectra stimulate each of the different photoreceptors in the retina. In birds, these photoreceptors are the four single cones used for colour vision, which are sensitive to long (L), medium (M), short (S), and very short (VS) wavelengths of light (Cuthill 2006).

To analyse chromatic variation, the quantum catches for each cone type were converted into three chromatic coordinates (x, y and z), where Euclidean distances between points reflect perceptual differences, using the formulas of Cassey *et al.* (2008). As there are no data for the song thrush, *Turdus philomelos*, which is the main predator of *Cepaea*, we used sensitivity functions for the closest available relative, the blackbird *Turdus merula* (Hart et al. 2000; Hart 2001), namely cone proportions of VS: 0.528, S: 0.904, M: 1.128, and L: 1, sensitivity functions of 373, 461, 543 and 603, respectively. The analysis assumed that the L cone has a noise-to-signal ratio of 0.05, so that the ratios for the other cones were VS: 0.0688, S: 0.0526 and M: 0.0471. The irradiance spectrum of ‘standard daylight’ (d65) was used for the main analyses. However, analyses were also run for ‘woodland shade’ to understand the influence of illuminant on avian perception of colour (Vorobyev et al. 1998).

To identify the main axes of chromatic variation, we carried out a Principal Components Analysis (PCA) on the chromatic coordinates (x, y and z), preserving the perceptual distances (JNDs) by using a covariance matrix rather than a correlation matrix (Delhey et al. 2015). To understand whether there are potential clusters within the chromatic coordinate data, Gaussian mixture modelling was carried out using Mclust 5.3 in R version 3.3.3 (Scrucca et al. 2016). A number of models were compared, each of which assumed a different number of clusters (from 1 to 10), normally distributed in multivariate chromatic space. Several classes of model were considered, each with a different assumption about the homogeneity of variance and orientation among clusters. The best fitting model was then determined as the one with highest Bayesian Information Criteria (BIC), with significant differences determined using a bootstrap approach.

The methods of Delhey et al. (2015) were also used to assess achromatic variation. In birds, sensitivity to achromatic cues is supposed to be mediated by double cones which have the same pigment as L cones in birds but different oil droplets, so have a wider sensitivity range. Values of achromatic contrast were therefore estimated, again in JNDs, by computing achromatic contrast between each reflectance spectrum and a reference (a very low value of double cone quantum catch, 0.001), corresponding to a dark spectrum, and using the same noise-to-signal ratio.

### Analysis of morph frequencies

We investigated evidence for effects of location and banding on the likelihood that a snail belonged to a particular morph, using generalised linear mixed effects models (GLMMs) with binomial errors. Each morph was considered separately, with each snail to be scored as belonging to the focal morph (1) or not (0). The three analyses are not independent, since each snail can only belong to one morph. Banding pattern was fitted as a fixed factor, whilst the effect of geographic location was examined at three spatial scales. Variation in morph frequency at a local level was modelled with random effect for site. Variation at a regional level was considered by fitting a fixed effect of geographic region. Finally, continental scale variation was modelled by looking for fixed linear and quadratic effects of latitude and longitude. The fact that region and latitude/longitude are partially collinear was reflected in the model-fitting procedure. We first fitted a saturated model with all main effects, except for region, and their two-way interactions (excluding interactions involving quadratic effects). Then, fixed terms were removed in a stepwise fashion, testing the effect of deletion using likelihood ratio tests, until only significant terms remained. Effects of latitude/longitude were then substituted with an effect of region and we compared the Akaike Information Criterion (AIC) of the resulting models, to test if region was better at capturing any large-scale geographic variation. Testing random effects in generalised linear mixed models is problematic, so we compared the AIC of the saturated GLMM with that of a generalised linear model without the random term for site to provide an approximate test of the importance of site.

## Results

### Variation in colour

We measured the individual reflectance spectra of 1172 shells, mainly collected from across Europe (Table S1; Figure 1) and then transformed them into visual space coordinates, xyz. To visualise this chromatic variation, and the relationship with human-scored and Mclust-defined colour categories (below), the xyz coordinates were plotted in visual colour space (Figure 2). There were no obviously discrete groups.

**Figure 2.**
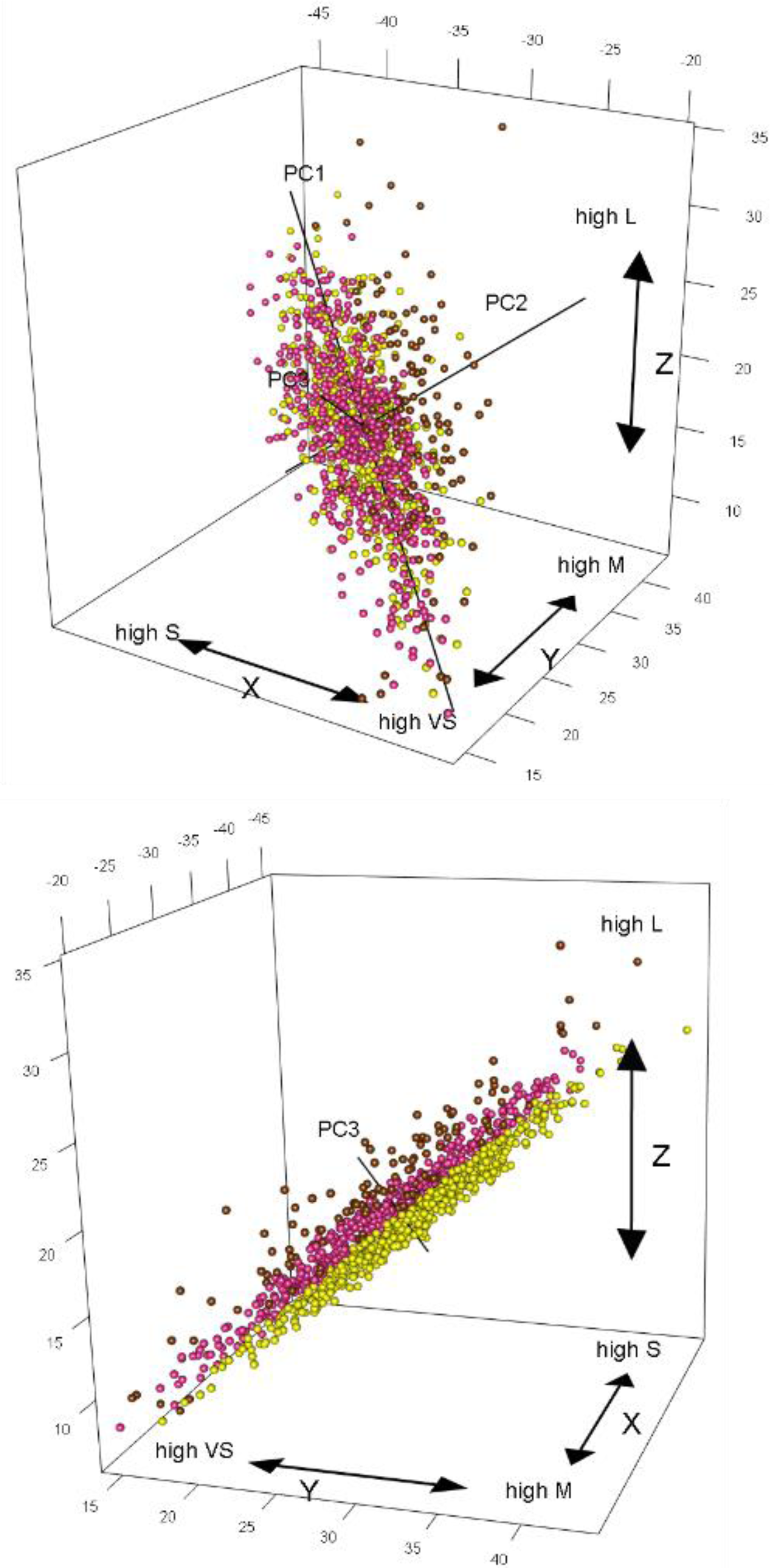
Axes of chromatic variation in the shell of *C. nemoralis*, using avian visual space, shown from two different perspectives (see also Supplementary Movie 2). Units on x, y and z axes are in JNDs. The solid lines illustrate variation along the first three principal components; individual points are coloured according to Mclust classification of the shell, either yellow, pink or brown. Top: Variation along PC1 (87%) mainly represents differences in saturation between shells. PC2 (11%) shows relatively higher stimulation of L cones and lesser stimulation of M and S cones, and tends to separate brown from pink/yellow. Bottom: PC3 (2%) shows relatively high stimulation of the M cones compared to lesser stimulation of the S and L cones, and tends to separate yellow from pink and pink from brown.

A PCA on the xyz coordinates showed a first axis which explained 87% of chromatic variation. PC1 had a moderate positive loading for x (0.61), and a moderate negative loading for y (−0.64) and z (−0.46). Two further axes explained 11% and 2% of the variation, the second having a positive loading on all axes (0.75, 0.28, 0.61, respectively), and the third a mixture (−0.26, −0.71, 0.65). The range of observed variation on each axis was considerable: 41, 22 and 8 JNDs for x, y and z respectively (Figure 2). Plotting the average normalized reflectance spectra for each quartile of each principal component showed how the three PC axes correspond to chromatic variation (Figure 3). Variation along PC1 represents relatively high stimulation of L cones and lesser stimulation of S cones, relative to M cones. Variation in PC2 showed relatively higher stimulation of L cones and lesser stimulation of M cones. PC3 showed relatively high stimulation of the M cones compared to lesser stimulation of the S and L cones. Only PC1 showed any differences in the VS region but the shells barely reflected in the UV.

**Figure 3.**
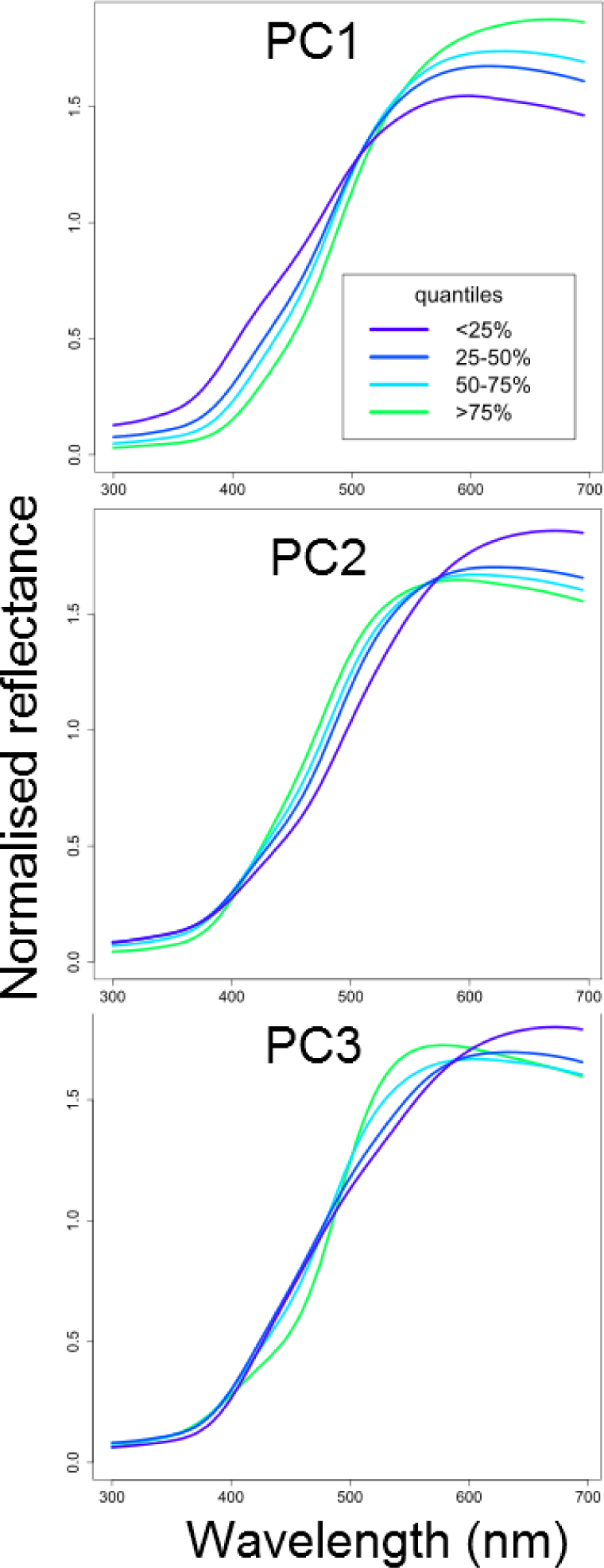
Interquartile ranges of the average normalised reflectance spectra for the principal component axes shown in Figure 2. These plots confirm that variation on PC1 mainly represents differences in saturation between shells; PC2 represents relatively higher stimulation of L cones and lesser stimulation of M and S cones; PC3 represents relatively high stimulation of the M cones compared to lesser stimulation of the S and L cones.

To investigate whether snail shells cluster in chromatic space, and whether observed clusters correspond to human-scored qualitative colour morphs, Gaussian finite mixture modelling was applied to the xyz visual space coordinates. The best model (VVV, ellipsoidal, varying volume, shape, and orientation; BIC −15727.3; *P* < 0.001 compared 2^nd^ best model) recovered three clusters, roughly corresponding to human-scored yellow (46%, n=539), pink (44%, n=511) and brown (10%, n=122) (Table 1). The next best fitting model also recovered three clusters (VEV; BIC - 15749.3; *P* < 0.001 compared with 3^rd^ best model) and the third recovered four clusters (EEV; BIC −15755.5; the 4^th^ cluster contained only 16 individuals).

**Table 1.**
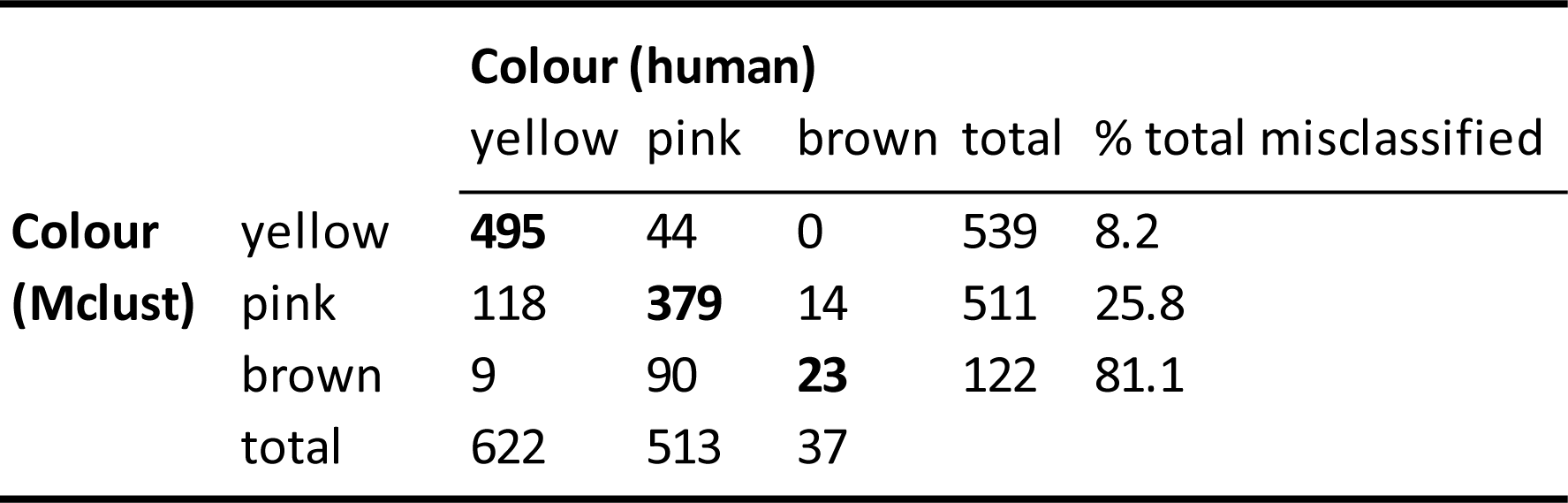
Comparison between human perceived colour categories and Mclust-defined groups. Shells that were scored the same are on the diagonal (in bold). Yellow and pink were most common, and so the absolute number of discordant scores was relatively low. Brown had by far the highest proportion of discordant scores.

Comparing human-scored (A.D.) and Mclust-defined groups, the overall concordance was good at 76% (Table 1), with a similar error rate (75%) for the student group. The highest proportion of discordant scores were human-scored yellow shells that Mclust classed as pink (10% for A.D.; 12% for student group), with the other major discrepancies being human-pink classed as Mclust-brown (8%), and human-yellow classed as Mclust-pink (4%). The main difference was that human-scoring reported relatively few brown shells (n=37), whereas the same group in Mclust was larger (n=122). With misclassifications adjusted relative to the total number of each Mclust shell type, 81% of the brown group were in a different human-scored group (74% pink, 7% yellow), compared to 26% of the pinks (3% brown 23% yellow) and just 8% of yellows (8% pink, 0% brown). Thus, while the overall correspondence between human and Mclust scoring of shell colour was good, the yellows were scored accurately (92%), pinks less so (74%) and brown poorly (19%).

Plots of human-scored colours along the three PCs (Supplementary Movie 1) and Mclust-categories were concordant with the above analyses (Figures 2, 4; Supplementary Movie 2). Broadly, PC1 did not separate different human-perceived colours or categories of shell, but instead mainly represents differences in saturation, or purity of colour, between individuals (Figure 4). PC2 separated brown from yellow and pink, and PC3 broadly separated all three colours, yellow, pink and brown.

**Figure 4.**
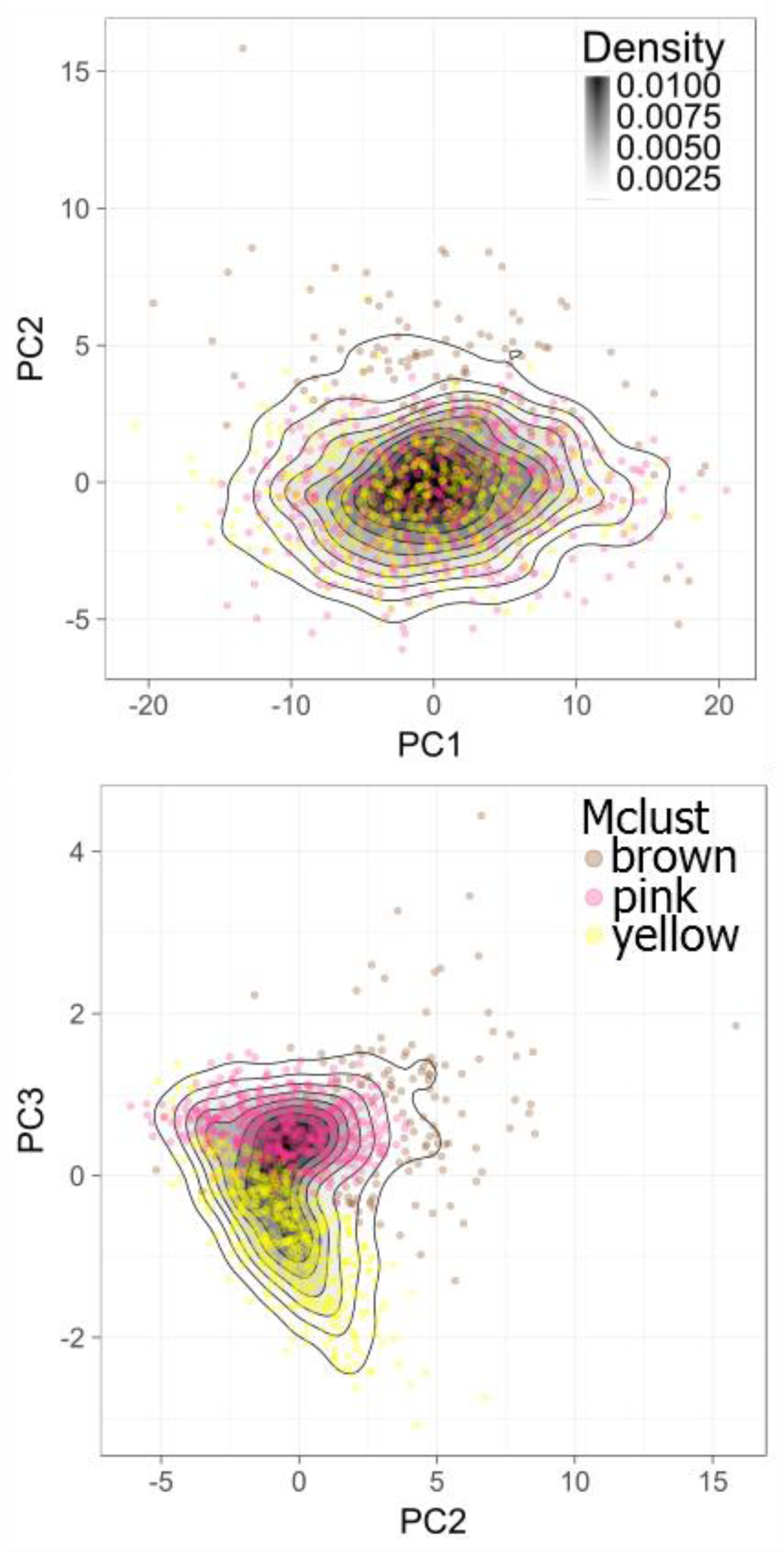
Scatterplot and associated density plot, showing variation along three principal component axes. Units are in JNDs. Points are coloured according to Mclust classification of the shell, either yellow, pink or brown. Top: PC1 versus PC2. Bottom: PC2 versus PC3.

The above analyses were repeated using woodland shade rather than standard daylight. The main difference was that while Mclust again recovered three groups, brown shells were more common (14%, n=168), with fewer pinks (40%, n=474) and approximately the same number of yellows (45%, n=530).

Finally, achromatic variation was also considerable, varying over more than 100 JNDs, and without any obvious differences between Mclust-defined colour morphs (Supplementary Figure 1).

### Geographic variation between morphs

Large-scale geographic variables (latitude, longitude and region) had significant effects on the probability that a snail was pink or yellow, but not the probability that a snail was brown (Table 2). Pink morphs were significantly less common at mid-latitudes (Figure 5). Snails with more than one band (B) and those which were mid-banded (M) were more likely to be pink in the west, while unbanded (O) snails were more likely to be pink in the east (Figure 5). In contrast, yellow snails were less common at high latitudes, and were affected by an interaction between longitude and banding which was the reciprocal of that seen in pink snails (Figure 6). Morph frequencies also varied at a local level: a saturated mixed model including the random effect of site was much better (Brown AIC = 443.7; Pink AIC = 868.9; Yellow AIC = 852.8) than an equivalent model without the random effect (Brown AIC = 646.4; Pink AIC = 1407.9; Yellow AIC = 1416.2). Banding was associated with colour morph in various ways. In addition to the interaction between banding and longitude in pink and yellow snails mentioned above, unbanded snails (O) were generally more likely to be brown (14% of all unbanded snails), than snails that were mid-banded (M; 11.6%) or had several bands (B; 5.8%).

**Table 2.**
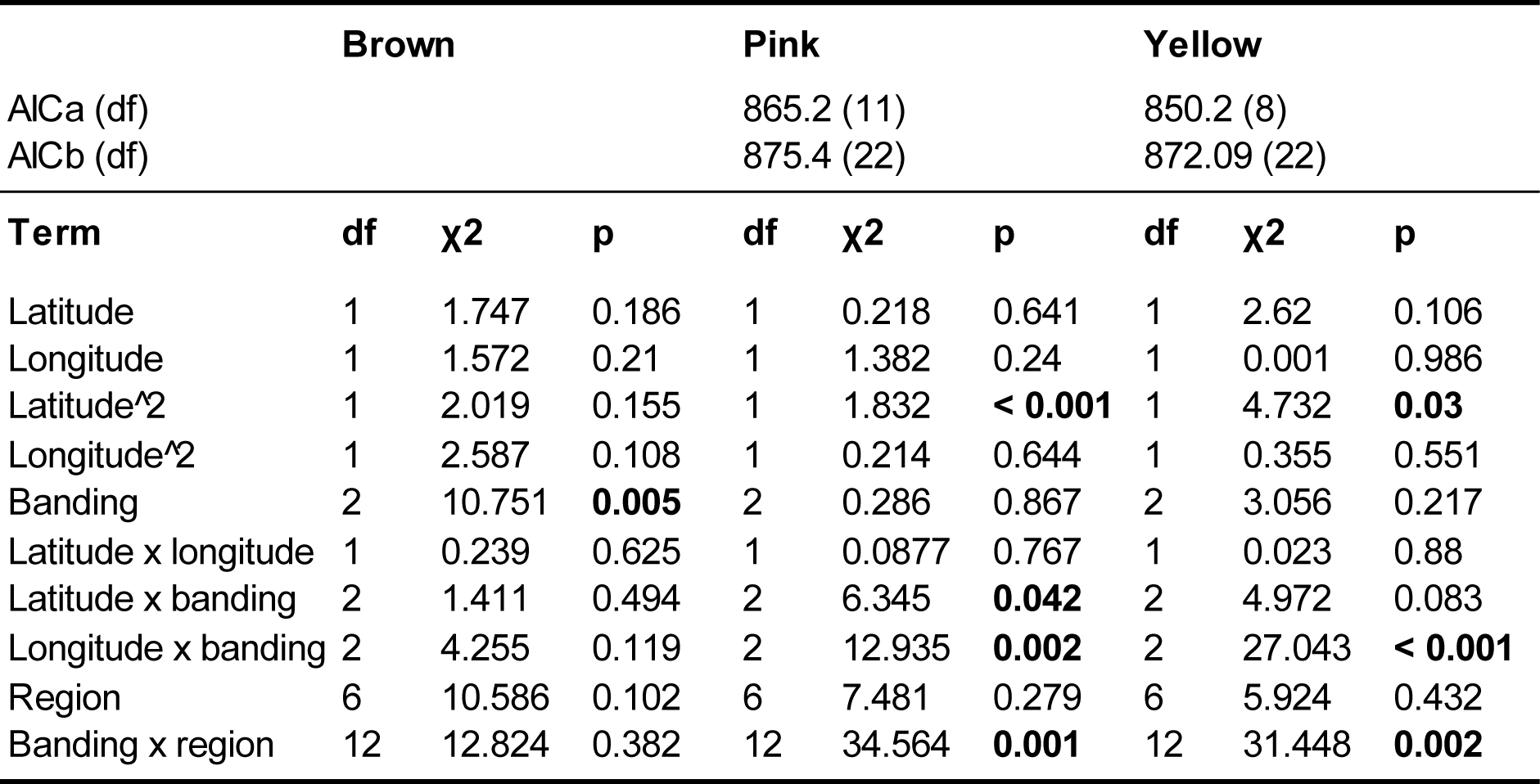
Results of likelihood ratio tests of the terms in binomial GLMMs of the effects of geographic variables and banding phenotype on the probability that a snail belonged to each of the three colour morphs. Significant p-values are in bold. The effects of modelling large-scale geographic variation in two ways are illustrated by the AIC values for the best model in which linear and quadratic effects of latitude and longitude were included (AICa), and the best model in which geographic region was included (AICb). All models include a random effect for site.

|  | **Brown** |  |  | **Pink** |  |  | **Yellow** |  |  |
| --- | --- | --- | --- | --- | --- | --- | --- | --- | --- |
| AICa (df) |  |  |  | 865.2 (11) |  |  | 850.2 (8) |  |  |
| AICb (df) |  |  |  | 875.4 (22) |  |  | 872.09 (22) |  |  |
| **Term** | **df** | **χ<sup>2</sup>** | **p** | **df** | **χ<sup>2</sup>** | **p** | **df** | **χ<sup>2</sup>** | **p** |
| Latitude | 1 | 1.747 | 0.186 | 1 | 0.218 | 0.641 | 1 | 2.62 | 0.106 |
| Longitude | 1 | 1.572 | 0.21 | 1 | 1.382 | 0.24 | 1 | 0.001 | 0.986 |
| Latitude<sup>2</sup> | 1 | 2.019 | 0.155 | 1 | 1.832 | **< 0.001** | 1 | 4.732 | **0.03** |
| Longitude<sup>2</sup> | 1 | 2.587 | 0.108 | 1 | 0.214 | 0.644 | 1 | 0.355 | 0.551 |
| Banding | 2 | 10.751 | **0.005** | 2 | 0.286 | 0.867 | 2 | 3.056 | 0.217 |
| Latitude x longitude | 1 | 0.239 | 0.625 | 1 | 0.0877 | 0.767 | 1 | 0.023 | 0.88 |
| Latitude x banding | 2 | 1.411 | 0.494 | 2 | 6.345 | **0.042** | 2 | 4.972 | 0.083 |
| Longitude x banding | 2 | 4.255 | 0.119 | 2 | 12.935 | **0.002** | 2 | 27.043 | **< 0.001** |
| Region | 6 | 10.586 | 0.102 | 6 | 7.481 | 0.279 | 6 | 5.924 | 0.432 |
| Banding x region | 12 | 12.824 | 0.382 | 12 | 34.564 | **0.001** | 12 | 31.448 | **0.002** |

**Figure 5.**
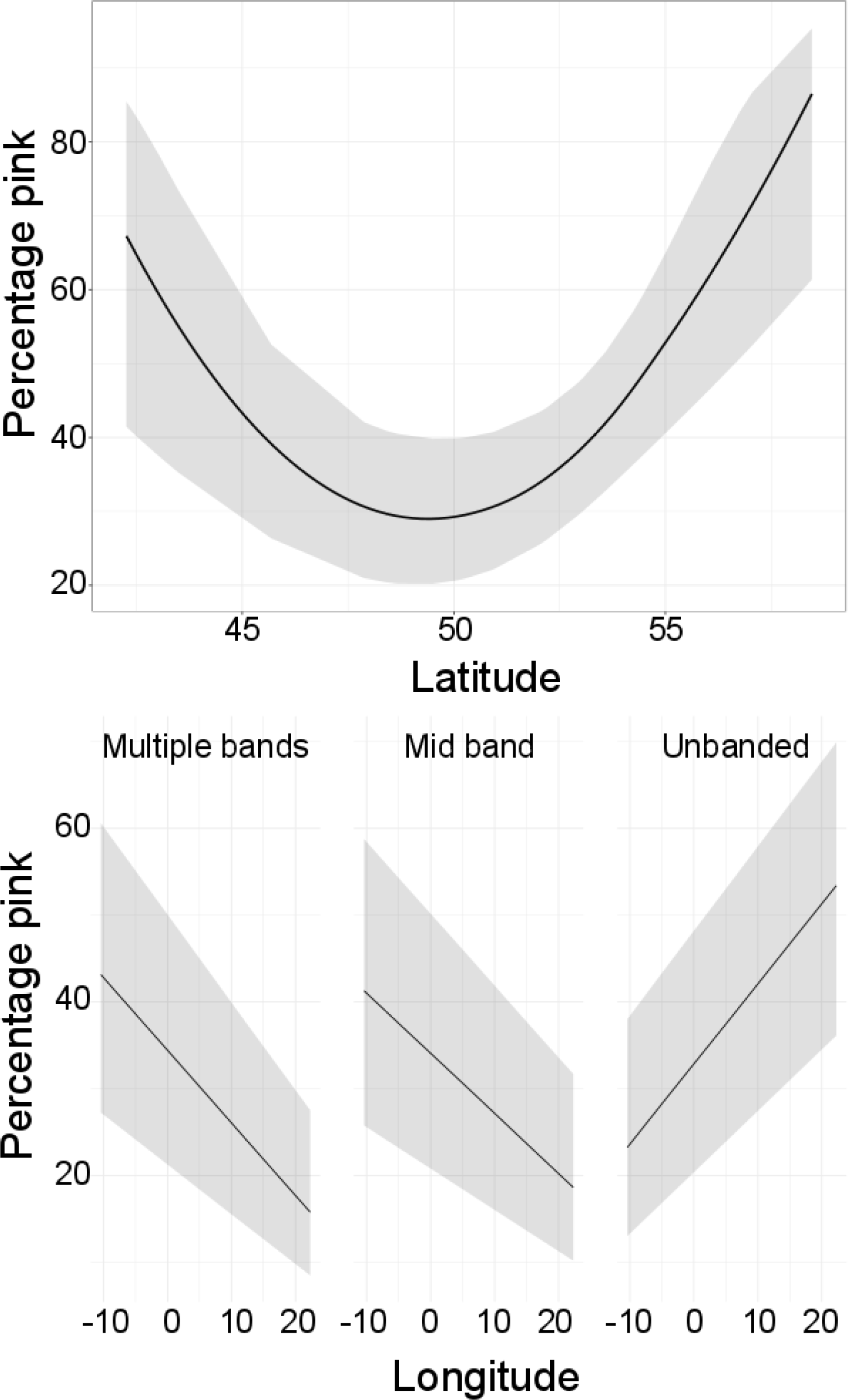
Scaled effects of latitude on the proportion of pink shells (top), and longitude on banding and proportion of pink shells.

**Figure 6.**
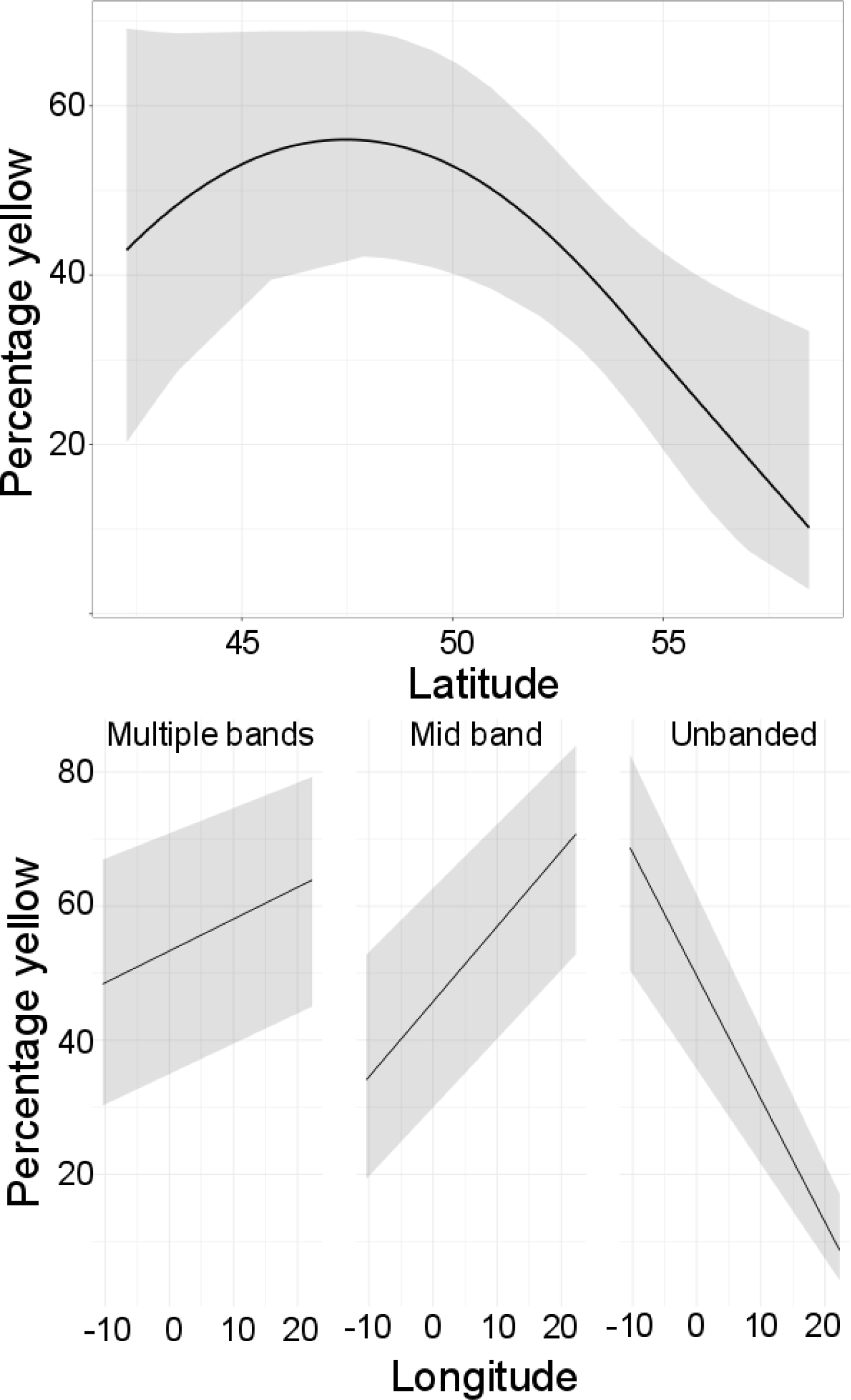
Scaled effects of latitude on the proportion of yellow shells (top), and longitude on banding and proportion of yellow shells.

## Discussion

By measuring the ground colour of *Cepaea nemoralis* shells collected across the breadth of the European distribution, we used psychophysical models of avian vision to understand how the shell colour may be perceived by birds, and to describe how this varies in geographic space, and with respect to other characters such as banding. The findings have significance for understanding the *Cepaea* polymorphism, and the nature of the selection that acts upon it, as well as more generally highlighting the need to objectively measure colour variation in other systems before beginning to test for possible explanations.

Broadly, we found that both chromatic (Figure 2) and achromatic variation (Supplementary Figure 1) is considerable, occurring over many perceptual units (JNDs). If this variation, both within and among human-perceived colour morphs, affects prey detection or identification by avian predators, then the presumption is that the polymorphism must be impacted by natural selection. The current available evidence suggests that animals in general use chromatic and achromatic signals for separate tasks, for example, using achromatic signals to identify the location, shape and motion of objects, while chromatic signals identify surface quality (Osorio and Vorobyev 2005). However, while this is also likely the case for avian predators, specific experimental evidence from birds is sparse (Osorio et al. 1999; Kang et al. 2015; White and Kemp 2016).

In our analysis, we found that chromatic variation in shells is continuously distributed in visual space, meaning that there are no wholly discrete colours (Figure 2). Perhaps surprisingly, we found that the most variable chromatic axis (PC1; 87%) that would be visible to a bird reflects the degree of saturation, or purity of colour. Axes separating human-perceived colours showed less variation, PC2 (11%) separating brown from yellow/pink, and PC3 (2%) broadly separating yellow, pink and brown.

Despite the lack of discrete colours, density-based clustering recovered three main shell types, which roughly correspond to human-perceived yellow, pink and brown (Table 1; Figure 3). Brown shells were more common according to the objective analysis than perceived by humans, with the frequency higher again when using woodland shade as an illuminant. Therefore, prior studies that (necessarily) used changes in frequencies of human-perceived colours to understand natural selection on snail shells may have missed a significant part of the picture – not only may birds use both achromatic and chromatic cues to differentiate morphs, but they should also be able to perceive chromatic differences to a much finer precision than a simple trivariate yellow, pink or brown categorisation that humans are obliged to use in qualitative surveys. Of course, this does not mean that birds react to the many morphs equally – it is possible that they categorically perceive a continuous variable (Caves et al. 2018). Further investigations are needed, especially using a bird such as the song thrush.

The effects of geographic location and banding pattern on variation in the reflectance spectrum of snails were also examined, the initial aim being to develop methods to *describe* variation, rather to *explain* it (e.g. by looking for correlations with environmental variables, putative selective agents, etc., as others have done; Silvertown et al. 2011). Generally, we found that geographic variables (latitude, longitude and region) and banding are generally associated with different frequencies of the three traditional colour morphs, with the main directional trend being that yellow snails are most common at mid-latitudes, as was found in much larger studies (Jones et al. 1977; Silvertown et al. 2011). Similarly, as previously reported (Cain et al. 1960), epistasis meant that unbanded snails (O) were generally more likely to be brown, and banded snails (B) were less likely to be brown. Therefore, by establishing a method for quantitatively measuring colour, and showing that a relatively small sample can be used to infer wide geographic patterns, this work provides a baseline for further studies on the polymorphism.

### Discrete or indiscrete?

Laboratory crosses in the past have revealed that the variation in the *Cepaea* shell phenotype is predominantly controlled by a ‘supergene’, which in a recent definition is a *genetic architecture involving multiple linked functional genetic elements that allows switching between discrete, complex phenotypes maintained in a stable local polymorphism* (Thompson and Jiggins 2014; Llaurens et al. 2017). This meaning fits with the traditional view – and the classical ‘Fordian’ theory of polymorphism (Ford 1964) – that the ground colour of the shell is one of three more or less discrete colour classes, either yellow, pink or brown, and indeed, is part of the reason that *Cepaea* snails became a well-studied system. However, while scoring the shell colour into different, discrete types is straightforward in offspring of individual crosses in the lab, the acknowledged reality is that it is sometimes difficult to classify shells consistently (e.g. see Table 1), especially when they are apparently intermediate in form.

Now, in our study, we have shown definitively that the colour polymorphism of *Cepaea nemoralis* is not discrete (Figures 2, 4). This finding emphasises the specific practical problem for projects collecting and using shell polymorphism data, especially those based entirely in the field and using citizen science (e.g. Evolution Megalab; Silvertown et al. 2011). However, it also illustrates a more general problem: if we do not have a precise definition of the *Cepaea* polymorphism and an understanding of the underlying genetic control, then how can we claim to understand the evolutionary and ecological factors that maintain colour variation?

Mathematical modelling is one method that can be used to explore the evolution of polymorphism, and of most relevance to this work, the circumstances that may or may not lead to discrete phenotypes. Historically, the argument of Ford was that despite the fact that supergenes may appear as Mendelian loci, they were actually rather complicated arrangements of several loci that are effectively prevented from being broken up by recombination under most normal circumstances. Thus, in both colour polymorphism in general (e.g. in side-blotched lizards; Sinervo and Lively 1996) and specifically relating to supergenes (e.g. in butterfly mimicry rings; Joron et al. 2011; Kunte et al. 2014), the distinctiveness of the morphs is a central feature of the genetic control; the genetic architecture specifically prevents phenotypes from “dissolving” into continuous trait distributions (Ford 1964).

Much of the existing research has therefore begun from the premise of understanding how evidently discrete types come about, and thus give insight into the adaptive evolution of genome structure (Cuthill et al. 2017). In simulations it has been have shown that natural selection tends to favour lowered recombination when intermediate genotypes are at a disadvantage; unlinked loci modify the phenotype to adapt to local conditions (e.g. to a local Batesian model butterfly; Charlesworth and Charlesworth 1975b, a; Llaurens et al. 2017). More recently, and perhaps most directly relevant to understanding the *Cepaea* polymorphism, Kopp and Hermisson (2006) devised a model to investigate the evolution of a quantitative trait under frequency-dependent disruptive selection. Their finding was that over generations most of the genetic variation tends to concentrate on a small number of loci.

The historic background is perhaps part of the reason that most of the recent progress in understanding supergenes has mainly come from species or systems that show simple, wholly discrete phenotypes, for example in butterfly mimicry rings (Joron et al. 2011), or heterostylous plants (Li et al. 2016). However, in *Cepaea* there are many colour morphs, such that colour variation is quantitative and due to a supergene; in other species such as the guppy *Poecilia* and the cichlid *Labeotropheus*, the inheritance of often considerable colour variation is due to several loci, some sex-linked and others not (Tripathi et al. 2009; Thompson and Jiggins 2014; Wellenreuther et al. 2014). Thus, developing theory on the impact of negative frequency-dependent (apostatic) selection must be able to account for these complexities, including those where supergenes are absent and variation is quantitative, otherwise there is a risk that models will simply reaffirm what we already know.

In one recent model, it was shown that crypsis and apostatic selection together may act to maintain a large number of morphs within a population, and in another apostatic selection was shown to maintain variation between similar species (Franks and Oxford 2011, 2017). In another more recent study, a simulation was used to explore the influence of predator perspective, selection, migration, and genetic linkage on colour allele frequencies. The relative sizes of predator and prey home ranges can result in large differences in morph composition between neighbouring populations (Holmes et al. 2017). Finally, in an empirical study blue jays *Cyanocitta cristata* searched for digital moths on mixtures of dark and light patches at different scales of heterogeneity. It was found that complex backgrounds with many moth-like features elicited a slow, serial search that depended heavily on selective attention. The result was increased apostatic selection, producing a broad range of moth phenotypes (Bond and Kamil 2006). All of these circumstances may apply to the *Cepaea* colour polymorphism.

Overall, there is an open debate – but little empirical data – on how the relative heterogeneity of the environment/substrate, density, distance or motion may influence the selection for crypsis or negative frequency dependence (Cuthill et al. 2017; Barnett et al. 2018). As Surmacki *et al.* (2013) summarised, if heterogeneous areas consist of large patches of diverse habitats then this may promote the evolution of specialist morphs through selection for crypsis, producing a few distinct or specialist morphs, each more or less well matched to the coloration of the preferred habitat type (Endler 1978; Bond 2007). If instead there are a mixture of small microhabitats, apostatic selection is more likely to result in multiple morphs that may be equally cryptic in all “grains” of the habitat. This is because in such circumstances, predators use search images of the most common morph, and this can lead to frequency-dependent selection.

### Supergenes return

In contrast to a relative paucity of field data, and a relatively lack of progress in establishing baseline theory, advances in DNA sequencing technology have meant that knowledge on the genetics of colour polymorphism is advancing rapidly. As hypothesised, in the still relatively few supergenes that have been fully characterised, the discrete phenotypes are maintained due to close physical proximity of the gene(s) and/or tight linkage (Joron et al. 2011; Kunte et al. 2014; Gautier et al. 2018).

In comparison, a few more general studies on colour polymorphism, rather than on supergenes specifically, have begun to reveal the extent of phenotypic variation, and whether discrete or indiscrete. For example, reflectance spectra have been used to show that even though humans perceived the colour variation in the eggs of African cuckoo finch *Anomalospiza imberbis* as falling into discrete categories, the variation was actually continuous (Spottiswoode and Stevens 2010, 2011). Similarly, tawny dragon lizard *Ctenophorus descresii* does have discrete colour morphs, but there is still considerable variation within each morph (Teasdale et al. 2013). Further quantitative studies in other lizards in which colour polymorphism has traditionally been treated as qualitative are also increasingly showing that there are significant overlaps in colour (Cote et al. 2008; Vercken et al. 2008; Paterson and Blouin-Demers 2017).

In *Cepaea nemoralis* snails, the colour and banding elements of the supergene have been mapped (Richards et al. 2013) but we remain ignorant of the underlying genetics and the precise nature of the selection that acts upon the polymorphism. For instance, models of supergene evolution require that intermediate phenotypes are disadvantaged – this makes sense with respect to Batesian mimics or distylous flowers – but in snails a rare intermediate might be at an advantage, due to apostatic selection. At the molecular level, one scenario is that the extreme and effectively continuous colour variation of the shells is due to a corresponding high number of colour alleles within the supergene. An alternative scenario is that there actually relatively few colour alleles, with much of the chromatic variation due to effects of other modifying loci (Charlesworth and Charlesworth 1975b). A final consideration is that while colour variation might be continuous across a grand geographic scale, if most local populations are founded by few individuals, then local variation might be discrete, which is all that matters from a selective point of view. This is more likely to be the case when both colour and banding are considered as the visible phenotype, especially since they are frequently in linkage disequilibrium (Cook 2017).

Overall, by establishing a method for quantitatively measuring colour, this work provides a baseline for further studies on the polymorphism, both from the perspective of understanding the nature of selection, and ultimately, also the genes involved. To reconcile and test competing theories with the empirical observations, the next steps must be to identify the component parts and evolutionary origins of the supergene in *C. nemoralis*, develop a model of frequency-dependent selection, and further understand how birds react to specific elements of the chromatic variation. When all of these findings are brought together, only then can we begin to understand the evolutionary and ecological factors that maintain this “problem with too many solutions.” Whatever the final outcome, there is no risk that *Cepaea* snails will be relegated to “other adaptive polymorphism” (Thompson and Jiggins 2014), especially because, as Jones *et al.* (1977) suggested, it is important to study organisms for which polymorphism may be explained by a variety of processes, precisely because they are more realistic.

## Supplementary material

Supplementary Movies 1, 2. Supplementary Figure 1. Supplementary Table 1.

**Supplementary Figure 1.**
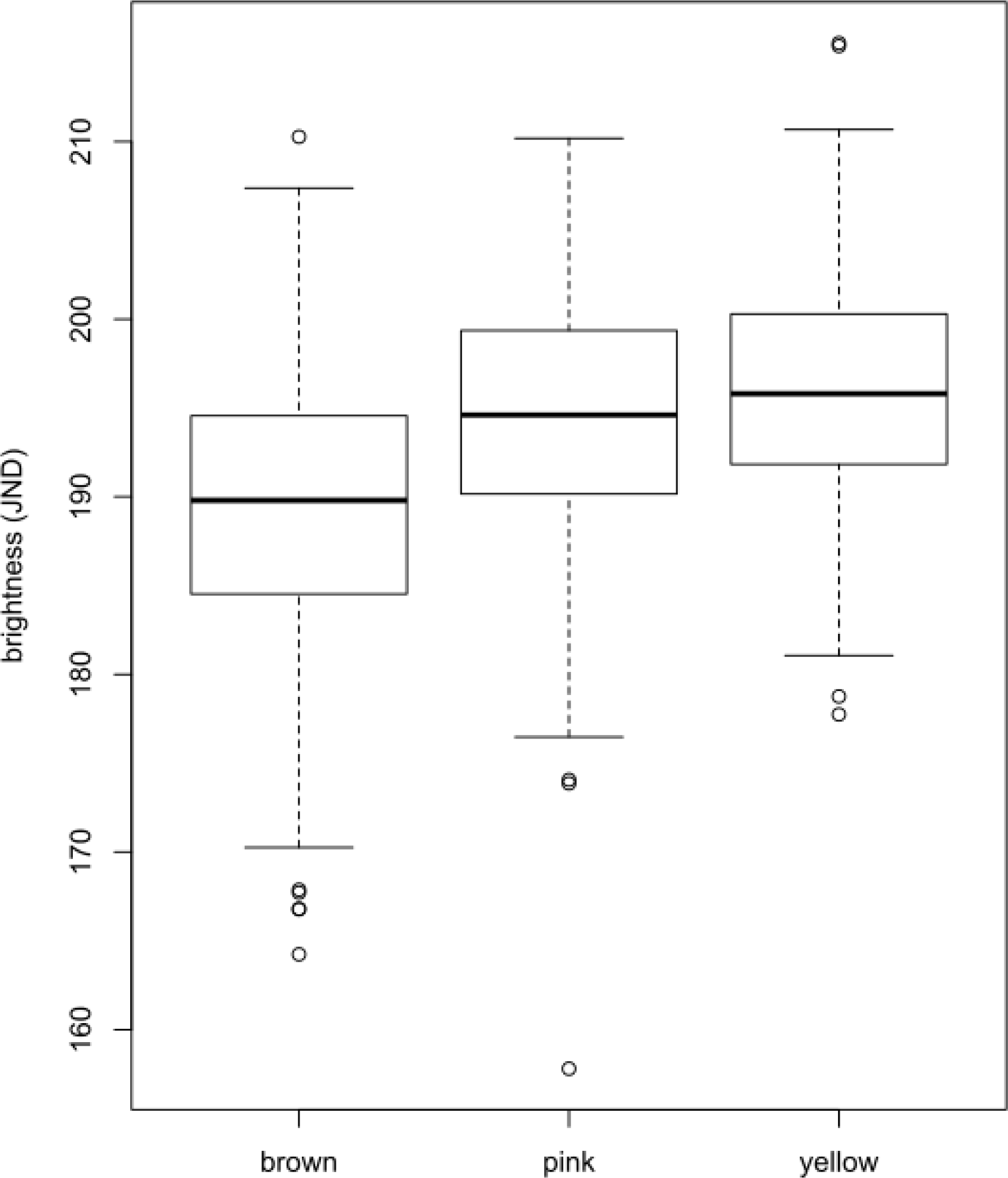
Boxplot showing extent of achromatic variation in Mclust-defined colour morphs of *Cepaea nemoralis*.

**Supplementary Movie 1.** Animation showing axes of chromatic variation in the shell of *C. nemoralis*, using avian visual space. Units on x, y and z axes are in JNDs. The solid lines illustrate variation along the first three principal components; individual points are coloured according to human-scoring of the shell, either yellow, pink or brown.

**Supplementary Movie 2.** Animation showing axes of chromatic variation in the shell of *C. nemoralis*, using avian visual space. Units on x, y and z axes are in JNDs. The solid lines illustrate variation along the first three principal components; individual points are coloured according to Mclust classification of the shell, either yellow, pink or brown.

## Acknowledgements

Thanks to both Alan Bond and Laurence Cook for helpful discussions, and to and Kaspar Delhey for discussion and assistance with the methods, to Alice Maiden and Shagufta Hadife for collecting data, and to Adele Grindon and a network of helpers collected the snails.

## Data archiving

Raw reflectance data will be included with the manuscript or uploaded to Dryad upon acceptance of the manuscript.

